# Distinct functions for the paralogous RBM41 and U11/U12-65K proteins in the minor spliceosome

**DOI:** 10.1101/2023.10.12.562036

**Authors:** Antto J. Norppa, Iftekhar Chowdhury, Laura E. van Rooijen, Janne J. Ravantti, Berend Snel, Markku Varjosalo, Mikko J. Frilander

## Abstract

In this work, we identify RBM41 as a novel unique protein component of the minor spliceosome. RBM41 has no previously recognized cellular function but has been identified as a paralog of the U11/U12-65K protein, a known unique component of the minor spliceosome that functions during the early steps of minor intron recognition as a component of the U11/U12 di-snRNP. We show that both proteins use their highly similar C-terminal RRMs to bind to 3’-terminal stem-loops in U12 and U6atac snRNAs with comparable affinity. Our BioID data indicate that the unique N-terminal domain of RBM41 is necessary for its association with complexes containing DHX8, an RNA helicase, which in the major spliceosome drives the release of mature mRNA from the spliceosome. Consistently, we show that RBM41 associates with excised U12-type intron lariats, is present in the U12 mono-snRNP, and is enriched in Cajal bodies, together suggesting that RBM41 functions in the post-splicing steps of the minor spliceosome assembly/disassembly cycle. This contrasts with the U11/U12-65K protein, which uses the N-terminal region to interact with U11 snRNP during the intron recognition step. Finally, we show that while RBM41 knockout cells are viable, they show alterations in the splicing of U12-type introns, particularly differential U12-type 3’ splice site usage. Together, our results highlight the role 3’-terminal stem-loop of U12 snRNA as a dynamic binding platform for the paralogous U11/U12-65K and RBM41 proteins, which function at distinct stages of minor spliceosome assembly/disassembly cycle.

## INTRODUCTION

In the majority of metazoan species, the removal of spliceosomal introns from pre-mRNA is carried out by two parallel ribonucleoprotein machineries: the major spliceosome, which splices the U2-type (major) introns, and the minor spliceosome, responsible for splicing of the U12-type (minor) introns. The overall architecture of the major and minor spliceosomes is similar. Both are composed of five small nuclear ribonucleoprotein (snRNP) particles—U1, U2, U4, U5 and U6 in the major spliceosome or U11, U12, U4atac, U5 and U6atac in the minor spliceosome—and additional non-snRNP splicing factors (Hall *et al*., 1996; Hastings *et al*., 2001; Kolossova *et al*., 1997; Tarn *et al*., 1996a, b; Yu *et al*., 1997). Compared to the major spliceosome, which has been extensively characterized structurally (Wilkinson *et al*., 2020), only a single high-resolution cryo-EM structure of a minor spliceosome activated for the first step of splicing (B^act^) (Bai *et al*., 2021) is presently available. However, it is generally accepted that the overall assembly pathway of the minor spliceosome is similar to that of the major spliceosome as both spliceosomes utilize similar snRNP complexes and a two-step splicing mechanism with branching and exon ligation reactions (Frilander *et al*., 2001; Tarn *et al*., 1996a, b). A key mechanistic difference is in the recognition of U12-type introns, which is carried out by a preformed U11/U12 di-snRNP that cooperatively binds to both the 5’ splice site (5’ss) and branch point sequence (BPS) (Frilander *et al*., 1999; Wassarman *et al*., 1992). In contrast to the earlier steps of the minor spliceosome assembly, very little is known about the post-catalytic events of the minor spliceosome cycle. In the major spliceosome, DHX8 (hPrp22) helicase drives the release of the ligated exon product from the post-catalytic (P) complex and turns it to the intron lariat spliceosome (ILS) which is subsequently disassembled by the DHX15 (hPrp43) helicase (Arenas *et al*., 1997; Company *et al*., 1991; Ohno *et al*., 1996). Recycling of snRNPs for subsequent rounds of splicing is thought to take place in the Cajal body, which is also the cellular site for other processes related to snRNP biogenesis and recycling (Staněk, 2017). There, the Cajal body-localized recycling factor SART3 is thought to function in both spliceosomes (Bell *et al*., 2002; Damianov *et al*., 2004).

Both the major and the minor spliceosome contain four unique small nuclear RNAs (snRNAs) and a number of unique protein components that are not found in the other spliceosome, while the majority of proteins are likely to be shared (Bai *et al*., 2021; Schneider *et al*., 2002; Will *et al*., 2004; Will *et al*., 1999). The first set of minor spliceosome-specific proteins (20K, 25K, 31K, 35K, 48K, 59K, 65K) were identified over 20 years ago by affinity purification and mass spectrometry analysis of the U11/U12 di-snRNP and U11 mono-snRNP fractions (Will *et al*., 2004; Will *et al*., 1999). Of these, U11/U12-65K, U11-59K and U11-48K form a chain of interactions connecting the U11 and U12 mono-snRNPs into a di-snRNP (Benecke *et al*., 2005; Turunen *et al*., 2008) and are essential for the stability of the di-snRNP (Argente *et al*., 2014; Norppa *et al*., 2018; Turunen *et al*., 2008). The U11 snRNP-associated U11-48K protein recognizes the 5’ splice site together with the U11 snRNA and interacts with the U11-59K protein (Turunen *et al*., 2008), which is further engaged in an interaction with the N-terminal part of U11/U12-65K (Benecke *et al*., 2005). The C-terminal RNA recognition motif (RRM) of U11/U12-65K binds the 3’-terminal stem-loop of U12 snRNA (Netter *et al*., 2009), but can also interact with the 3’-terminal stem-loop of the U6atac snRNA (Singh *et al*., 2016). ZRSR2, a component of the U11/U12 di-snRNP responsible for 3’ splice site recognition (Will *et al*., 2004), has also shown to almost exclusively affect splicing of U12-type introns in the cell (Madan *et al*., 2015), even though it may also have a separate role in the major spliceosome (Shen *et al*., 2010).

The notion that specific protein components of the minor spliceosome are needed only during the intron recognition phase and not in the later assembly steps has been challenged only very recently. The first such factor, the plant ortholog of RBM48 was originally identified as a U12-type intron splicing factor from a transposon screen in maize (Bai *et al*., 2019), and its specificity for U12-type introns was later confirmed in human cells (Siebert *et al*., 2022). A cryo-EM structure of the catalytically activated minor spliceosome (B^act^ complex) revealed that RBM48, together with additional three unique proteins ARMC7, SCNM1 and CRIPT, are all specific components of the B^act^ complex (Bai *et al*., 2021). The first unique protein component of the U4atac/U6atac di-snRNP and U4atac/U6atac.U5 tri-snRNP, CENATAC, was initially identified as a human disease gene and was shown to be specifically required for the splicing of U12-type introns with AT-AN termini (De Wolf *et al*., 2021). Similarly, mutations in DROL1, the plant ortholog of TXNL4B, the main interactor of CENATAC (De Wolf *et al*., 2021), were shown to lead to splicing defects with U12-type introns with AT-AC termini in *Arabidopsis thaliana* (Suzuki *et al*., 2022).

In this work, we provide evidence that the specific protein components in the minor spliceosome are not limited to intron recognition and catalytic steps. We show that RBM41, a closely related paralog of the U11/U12-65K, is a novel specific component of the minor spliceosome. The two paralogous proteins have a similar shared dual RNA binding specificity *in vitro*, interacting with the same 3’-terminal stem-loops of U12 and U6atac snRNAs with approximately equal affinity. However, while the U11/U12-65K functions during the intron recognition step, our results provide strong evidence that RBM41 functions in post-catalytic steps of the U12-type intron splicing. This further suggests that the 3’-terminal stem-loop of the U12 snRNA has a role as a protein-binding platform that is dynamically recognized first by the U11/U12-65K during the early steps of the spliceosome assembly followed by an exchange to RBM41 during or after the catalytic steps of splicing.

## MATERIALS AND METHODS

### Antibodies

The following antibodies were used in this study: anti-HA (BioLegend, 16B12), anti-GAPDH (Proteintech, 60004-1), anti-PDCD7 (Proteintech, 12485-1-AP), anti-RBM41 (Atlas Antibodies, HPA042881), anti-RNPC3 (Santa Cruz, sc-514951), anti-Sm (Invitrogen, MA5-13449), anti-SNRNP48 (Proteintech, 24297-1-AP), anti-V5 (Invitrogen, R960-25), anti-ZCRB1 (Bethyl Laboratories, A304-697A), Goat anti-Rabbit Alexa Fluor 488 (Thermo Fischer, A11008), Goat anti-Mouse Alexa Fluor 568 (Abcam, Ab175473), Mouse IgG2a kappa Isotype Control (Invitrogen, 14-4724-82). eBiocience Avidin-HRP (Invitrogen, 18-4100) was used to detect biotinylated proteins.

### Plasmid construction

RBM41 cDNA sequence, corresponding to Ensembl transcript ENST00000372479.7, was amplified from HEK293 cDNA in two fragments. Gibson assembly was used to assemble the fragments into vector pCI-neo in-frame with an N-terminal V5 epitope tag, resulting in pCI-neo-V5-RBM41. For BioID cell line construction, full-length RBM41 (1–413), RBM41 N-terminal fragment (1–258), RBM41 C-terminal fragment (259–413) and full-length 65K were cloned into MAC-tag-N vector (Addgene #108078, a gift from Markku Varjosalo) using Gateway cloning as described (Liu *et al*., 2020). Human DHX8 cDNA (MGC clone 5529639) was obtained from the Genome Biology Unit at the University of Helsinki and cloned into pCI-neo in-frame with an N-terminal V5 tag. Point mutations were introduced into plasmids using site-directed mutagenesis with Phusion polymerase.

### Expression and purification of recombinant proteins

C-terminal fragments of RBM41 (amino acids 267-413) and U11/U12-65K (amino acids 380-517) were expressed as GST-tagged proteins in *E. coli* Rosetta cells. Protein expression was induced with 0.5 mM IPTG for 3 h at 37°C and cells were harvested by centrifugation. Cell pellets were resuspended in 50 mM Tris-HCl (pH 7.5), 150 mM NaCl, 5 mM DTT, 1x cOmplete protease inhibitor cocktail (Roche) and lysed by sonication (30 s ON / 30 s OFF, 5 min total sonication time, Sonopuls HD 2070 with MS 73 microtip). After removal of cell debris (25,000 g, 15 min), GST-tagged proteins were captured from the lysate by incubation with glutathione agarose (Pierce) for 1 h at 4°C. The resin was washed 3 times with 20 volumes of Wash/Cleavage buffer (50 mM Tris-HCl (pH 7.5), 150 mM NaCl, 1 mM EDTA, 1 mM DTT) and GST tag cleaved by overnight digestion with PreScission Protease (Cytiva). Purified proteins were dialyzed against EMSA buffer (20 mM HEPES-KOH, 100 mM KCl, 1.5 mM MgCl2, 5% glycerol, pH 8.0) and protein concentrations measured by Pierce BCA protein assay (Thermo Scientific).

### Electrophoretic Mobility Shift Assay

Recombinant proteins (0.25–25 µM) were incubated for 1 h on ice with 5 nM [γ-^32^P]-ATP-labeled RNA oligonucleotides in binding buffer (20 mM HEPES–KOH (pH 8.0), 100 mM KCl, 1.5 mM MgCl_2_, 5% glycerol, 0.1 µg/µL BSA, 1 µg/µL yeast RNA (Roche), 1 U/µL RiboLock RNase inhibitor (Thermo Scientific)) in 10 µl final volume. After addition of 2.5 µl of 5x loading buffer (20 mM HEPES–KOH, 100 mM KCl, 1.5 mM MgCl2, 50% glycerol, 0.1% bromophenol blue, 0.1% xylene cyanol), 4 µl of each binding reaction was loaded onto a native polyacrylamide gel (6%, 80:1 acrylamide:bis-acrylamide ratio, 0.5x TBE, 5% glycerol). Electrophoresis was carried out at 120 V for 1.5 h at 4°C. After drying, the gel was exposed on an imaging plate and scanned using the FLA-5100 phosphorimager. Bound and free RNA bands were quantified using AIDA software (Raytest). Dissociation constants were determined by nonlinear regression using GraphPad Prism (One site – Specific binding) from three independent replicates.

### Cell culture and transfection

HEK293 cells were grown in DMEM, 10% FBS, 1% penicillin–streptomycin and 2 mM L-glutamine. All plasmid transfections were carried out using Lipofectamine 2000 (Thermo Fisher) according to manufacturer’s instructions.

### CRISPR/Cas9-mediated knockout of RBM41

HEK293 cells were transfected with pSpCas9(BB)-2A-Puro vectors with sgRNA sequences targeting RBM41 exon 2 or 3. 24 h after transfection, puromycin was added at 3 µg/ml to enrich for transfected cells and puromycin treatment continued for 72 h. After puromycin selection, monoclonal cell lines were obtained by limiting dilution in 96-well plates. Genomic DNA was extracted from single-cell clones using the NucleoSpin Tissue kit (Macherey– Nagel), and the targeted areas amplified by PCR using primers in the introns flanking exons 2 and 3 (Supplementary Table 1). Clones were screened for editing using the Surveyor Mutation Detection Kit (IDT). Positive clones were verified by TOPO cloning of PCR products followed by sequencing, as well as by direct sequencing of the PCR products and deconvolution of sequencing traces using the DECODR tool (Bloh *et al*., 2021).

### Northern blot

Northern blotting was carried out exactly as described (Norppa *et al*., 2021) using LNA or DNA oligonucleotide probes listed in Supplementary Table 1.

### RNA immunoprecipitation

For RNA immunoprecipitation with V5-tagged proteins, pCI-neo vectors for expressing V5-tagged proteins or empty pCI-neo vector (4 µg) were transfected into HEK293 cells in 6-well plate format using Lipofectamine 2000. For RNA immunoprecipitation with endogenous proteins in cell lysate, typically ∼20−10^6^ cells were used for one IP. 24 h after transfection, cells were washed with 1xPBS, scraped into lysis buffer (20 mM HEPES, pH 7.9, 137 mM NaCl, 10% glycerol, 1% NP-40 + 1x cOmplete protease inhibitor cocktail + 0.5 U/µl RiboLock) and sonicated (5×30 s, Bioruptor Twin, High setting). Cell debris was removed by centrifugation at 16,000 g for 15 min. The supernatant was incubated o/n at with rotation at 4°C with 2 µg IP antibody and for 1 h with Dynabeads Protein G (Invitrogen). Alternatively, the antibody was prebound to Dynabeads and incubated with lysate for 1 h at 4°C. After five washes with 200 µl lysis buffer, co-immunoprecipitated RNA was isolated by treatment of beads with Proteinase K, phenol:chloroform:isoamyl alcohol (pH 4.8) extraction and ethanol precipitation. For nuclear extract RIP experiments, 60 µl HeLa S3 nuclear extract was used for one IP. Nuclear extract was preincubated for 10 min at 37°C in 13 mM HEPES (pH 7.9), 2.4 mM MgCl_2_, 40 mM KCl, 2 mM DTT, 20 mM creatine phosphate, 0.5 mM ATP. After preincubation, the nuclear extract was diluted 1:5 and adjusted to 20 mM HEPES (pH 7.9), 137 mM KCl, 10% glycerol, 0.1% NP-40, 0.5 U/µl RiboLock, 1x cOmplete protease inhibitor. The remaining steps were carried out as described above.

### Glycerol gradient centrifugation

HeLa S3 nuclear extract (200 µl), prepared essentially as described in Tarn *et al*. (1994), was adjusted to 13 mM HEPES (pH 7.9), 2.4 mM MgCl_2_, 40 mM KCl, 2 mM DTT, 20 mM creatine phosphate, 0.5 mM ATP, and pre-incubated for 10 min at 30°C in a final volume of 300 µl. After incubation, 210 µl of 1x Gradient buffer (20 mM HEPES (pH 7.9), 40 mM KCl, 2 mM DTT, 2.4 mM MgCl_2_) was added, samples were centrifuged briefly (20,000 g, 1 min) and the supernatant was loaded on top of 10–30% glycerol gradient in 1x Gradient buffer. Gradients were centrifuged in a Sorvall TH-641 rotor at 29,000 rpm, 4°C for 18 h and fractionated using BioComp Piston Gradient Fractionator. Gradient preparation, centrifugation and fractionation was carried out by the HiLIFE Biocomplex unit at the University of Helsinki. For RNA extraction, 20% of each fraction was treated with Proteinase K, phenol:chloroform extracted and precipitated with ethanol. The remaining 80% of each fraction was precipitated with TCA for western blot analysis.

### BioID analysis

For each cell line, Flp-In™ T-REx™ 293 cells were grown in 5×15 cm plates to ∼70% confluency. MAC-tagged protein expression and biotinylation was induced by addition of 2 μg/ml of tetracycline and 50 μM biotin. Cells were harvested 24 h after induction by pipetting up and down with PBS-EDTA, centrifugation at 1,200 g for 5 min, and snap freezing the pellet in liquid nitrogen. Three independent replicates of 5×15 cm dishes were prepared for each cell line. BioID analysis was carried out essentially as described previously (Liu *et al*., 2018; Liu *et al*., 2020).

### cDNA synthesis and RT-PCR

RNA was treated with RQ1 RNase-free DNase (Promega) to remove any genomic DNA contamination. cDNA synthesis was carried out using Maxima H Minus RT (Thermo) and random primers according to the manufacturer’s instructions. For PCR, Phire polymerase (Thermo) and primers listed in Supplementary Table 1 were used.

### High-throughput sequencing

Total RNA isolated using Trizol extraction followed by an additional acidic phenol (pH 5.0) extraction. RNAseq libraries were constructed using Illumina TruSeq Stranded Total RNA kit (Illumina) Human Ribo-Zero rRNA depletion kit (Illumina). Paired-end 150+150 bp sequencing was performed at the Institute for Molecular Medicine Finland FIMM Genomics unit with Illumina NovaSeq 6000 using partial S4 flow cell lane. The STAR aligner (Dobin *et al*., 2012) was used for mapping the paired sequence reads to the genome (hg38/GRCh38). Transcript annotations were obtained from GENCODE (v29). The length of the genomic sequence flanking the annotated junctions (sjdbOverhang parameter) was set to 161. The Illumina adapter sequences AGATCGGAAGAGCACACGTCTGAACTCCAGTCAC and AGATCGGAAGAGCGTCGTGTAGGGAAAGAGTGTAGATCTCGGTGGTCGCCGTAT CATT were, respectively, clipped from the 3’ of the first and the second pairs in the read libraries (using clip3pAdapterSeq parameter). Statistics of the RNAseq data are presented in the Supplementary Table 5.

### Differential alternative splicing analysis and intron retention analysis

Differential alternative splicing (AS) analysis was done using Whippet (v0.11) (Sterne-Weiler *et al*., 2018). Both merged aligned reads (bam files) and AS event annotations from GENCODE (v29) were used to build the index reference for AS events. To detect the significantly differential events, probability cutoff of Pr > 0.9 and Percentage Spliced In deviation cutoff of |ΔΨ| > 0.05 were used. Differential intron retention was analyzed with IRFinder-S using SUPPA2 wrapper (Lorenzi *et al*., 2021; Middleton *et al*., 2017). A custom list of human U12-type intron coordinates (Supplementary Table 8) combining high-confidence U12-type introns from IntEREst (Oghabian *et al*., 2018), IAOD (Moyer *et al*., 2020), and MIDB (Olthof *et al*., 2019) databases was used in the annotation of U12-type introns and their host genes.

### Phylogenetic profiling-based co-evolution analysis

To conduct the phylogenetic profiling analysis of RBM41, we utilized a diverse set of eukaryotic proteomes. This dataset, consisting of 167 eukaryotic species, was previously compiled to represent the eukaryotic tree of life, and the species were selected based on their representation in the tree. We used automatic orthologous groups (OG) based on previous work (Deutekom *et al*., 2021; Vosseberg *et al*., 2021), generated using methods such as Orthofinder (Emms *et al*., 2019), eggNOG (Huerta-Cepas *et al*., 2019) hmm profile database, and OGs from Vosseberg *et al*. (2021). However, as RBM41 was not accurately represented in these automatically generated OGs, we manually created the OG for RBM41. This was achieved by performing a blast search with RBM41 against our in-house eukaryotic dataset and creating a Hidden Markov model (HMM) of RBM41, which was then used to perform a Hmm search (Mistry *et al*., 2013) against the same dataset. To determine the OG, a phylogenetic analysis was carried out with the top 100 entries of the Hmmer search using mafft E-INS-I (Katoh *et al*., 2005) and IQtree (Nguyen *et al*., 2015). In this phylogeny, a cluster with representatives from diverse eukaryotic groups indicated the RBM41 Orthologous group. Besides the support-value, the species overlap of this cluster with the other putative orthologous groups in the tree solidifies it as resulting from an ancient duplication and having been present in LECA. To compare the phylogenetic similarity, we computed cosine distances between the phylogenetic profile of each automatically generated OG and RBM41. As we suspected that RBM41 was part of the minor spliceosome, we also compared the phylogenetic profile of RBM41 to the profiles of other known spliceosome proteins obtained from Vosseberg *et al*. (2023).

## RESULTS

### RBM41 is a paralog of the U11/U12-65K protein and a putative component of the minor spliceosome

Human RBM41 is a ∼47 kDa protein with a single annotated domain, an RNA recognition motif (RRM) near the C-terminus of the protein (at positions 309–387) (Figure 1A). While no specific cellular function has been assigned to RBM41, it has been listed as a paralog of the U11/U12-65K protein (*RNPC3*), a structural component of the U11/U12 di-snRNP complex in the minor spliceosome (Benecke *et al*., 2005; Vosseberg *et al*., 2023). The paralog assignment is based on the local sequence similarity between the C-terminal regions of the two proteins, that encompass the core RRM and its N-terminal expansion (Figure 1B, 1C) which in the U11/U12-65K is essential for the stability of the C-terminal RRM and its interaction with the U12 snRNA (Netter *et al*., 2009). The RRM sequences of 65K and RBM41 conform with the general RRM consensus (Muto *et al*., 2012) except for an aromatic to nonaromatic amino acid substitution F/Y352Q at position 3 of the RNP1 motif (Figure 1B). The same substitution is present of in the homologous N-terminal RRMs of the U1A/U2B″/SNF family of spliceosome components, which are characterized by a YQF triad of RNP2 tyrosine and RNP1 glutamine and phenylalanine (Dekoster *et al*., 2014; Weber *et al*., 2018; Supplementary Figure 1). These three residues are displayed on the β-sheet surface of the RRM and engage in stacking interactions with RNA nucleobases (Dekoster *et al*., 2014; Weber *et al*., 2018). Conservation of the corresponding residues in the RBM41 RRM (Y312/Q352/F354; Supplementary Figure 1) suggests that RBM41 may employ a similar mode of RNA binding as utilized by the U1A/U2B″/SNF proteins.

**Figure 1.**
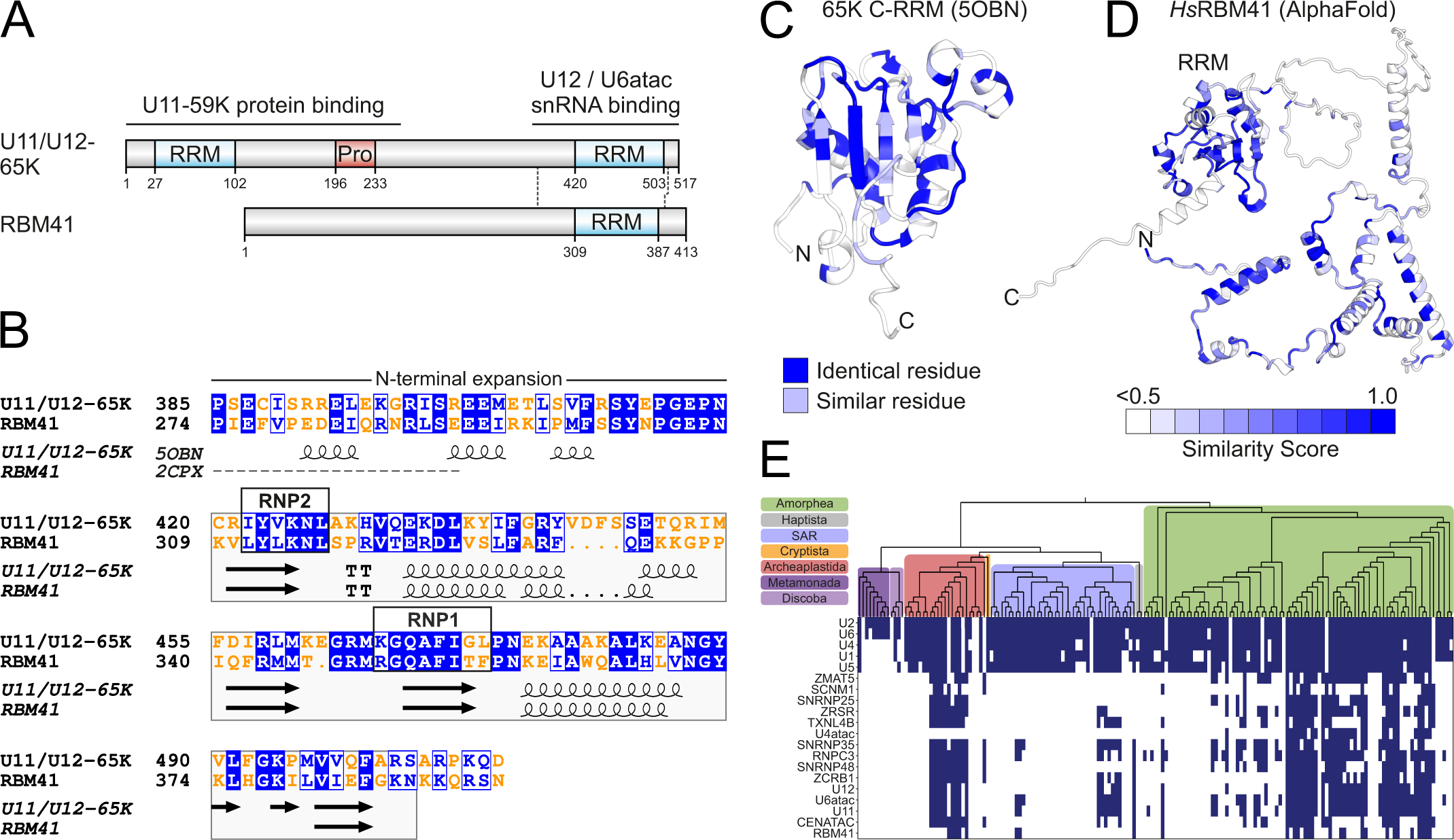
RBM41 is a paralog of the U11/U12-65K protein. (A) Domain structures of human U11/U12-65K and RBM41 proteins. (B) Pairwise sequence alignment of RBM41 and U11/U12-65K. Local sequence alignment was carried out using Matcher and visualized using ESPript 3.0 (Robert *et al*., 2014). Identical residues are shown in white text with blue background and similar residues in blue text with white background. Protein secondary structure elements extracted from NMR structures (U11/U12-65K: 5OBN, RBM41: 2CPX) are shown below the alignment. (C) Structure of the U11/U12-65K C-terminal RRM (5OBN) showing identical and similar residues between RBM41 and U11/U12-65K. (D) AlphaFold-predicted structure of human RBM41 colored for sequence conservation. Conservation is based on a multiple sequence alignment of RBM41 orthologues from 15 animal species (Supplementary Figure 2). Conservation was mapped to the structure with ESPript 3.0 and structure rendered using PyMOL. (E) Phylogenetic profile of RBM41 compared to the known minor spliceosome-specific proteins and minor and major spliceosomal snRNAs.

The homology between RBM41 and U11/U12-65K raises the question of whether RBM41 also functions in the minor spliceosome. We carried out a phylogenetic profiling-based co-evolution analysis in 167 eukaryotic species to identify proteins that show a similar phylogenetic presence/absence profile as RBM41 and may therefore function in the same molecular process (Figure 1E). Notably, two minor spliceosome-specific proteins, CENATAC and U11-48K (*SNRP48*) were found among the proteins showing the strongest co-occurrence with RBM41 (Supplementary Table 3), suggesting a role in the minor spliceosome. This was further supported by earlier reciprocal coevolution analysis for CENATAC (De Wolf *et al*., 2021), which similarly identified RBM41 as one of the top co-occurring proteins among the known minor spliceosome components. Upon comparing RBM41’s phylogenetic profile with that of other known minor spliceosome proteins (Figure 1E), it becomes evident that RBM41 adheres to the typical presence/absence profile of these proteins. However, the presence of RBM41 appears to be even less common than that of other minor spliceosome proteins.

In contrast to the homologous C-terminal RRMs, the N-termini of the two proteins do not share sequence similarity. This suggests a functional difference between RBM41 and U11/U12-65K, that uses the N-terminus to promote the formation of the U11/U12 di-snRNP via an interaction with the U11-59K protein (Benecke *et al*., 2005; Figure 1A). RBM41 N-terminus is highly conserved among animal orthologs (Supplementary Figure 2, Supplementary Table 2, Figure 1D) but lacks annotated domains and adopts a predominantly helical conformation in an AlphaFold prediction (Varadi *et al*., 2022) (Figure 1D). Together the sequence analysis suggests similar RNA binding properties, but otherwise divergent functions for the two paralogous proteins.

### RBM41 RRM interacts with the U12 and U6atac snRNAs in vitro

Previously, two independent systematic high-throughput studies of RNA-binding protein specificities (Dominguez *et al*., 2018; Ray *et al*., 2013) reported nearly identical consensus RNA motifs (Figure 2A) which are preferentially located in a loop sequence within an RNA stem-loop context. These RNA motifs bear a striking similarity to the loop sequences of the U12 snRNA 3’-terminal stem-loop and U6atac snRNA 3’-terminal stem-loop both of which are bound by the U11/U12-65K protein (Figure 2A, 2B; Benecke *et al*., 2005; Singh *et al*., 2016). Given the apparent RNA-binding similarities between the RBM41 and U11/U12-65K proteins, we compared their RNA-binding characteristics *in vitro,* using recombinant C-terminal RRMs containing additional N- and C-terminal regions known to be essential for RNA binding (Netter *et al*., 2009; Norppa *et al*., 2018). RNA binding properties were analysed using electrophoretic mobility shift assay (EMSA) with untagged RRMs and RNA oligonucleotides corresponding to the apical hairpins of the U12 snRNA and U6atac 3’-terminal stem-loops (Figure 2C, 2D; Benecke *et al*., 2005; Netter *et al*., 2009; Norppa *et al*., 2018).

**Figure 2.**
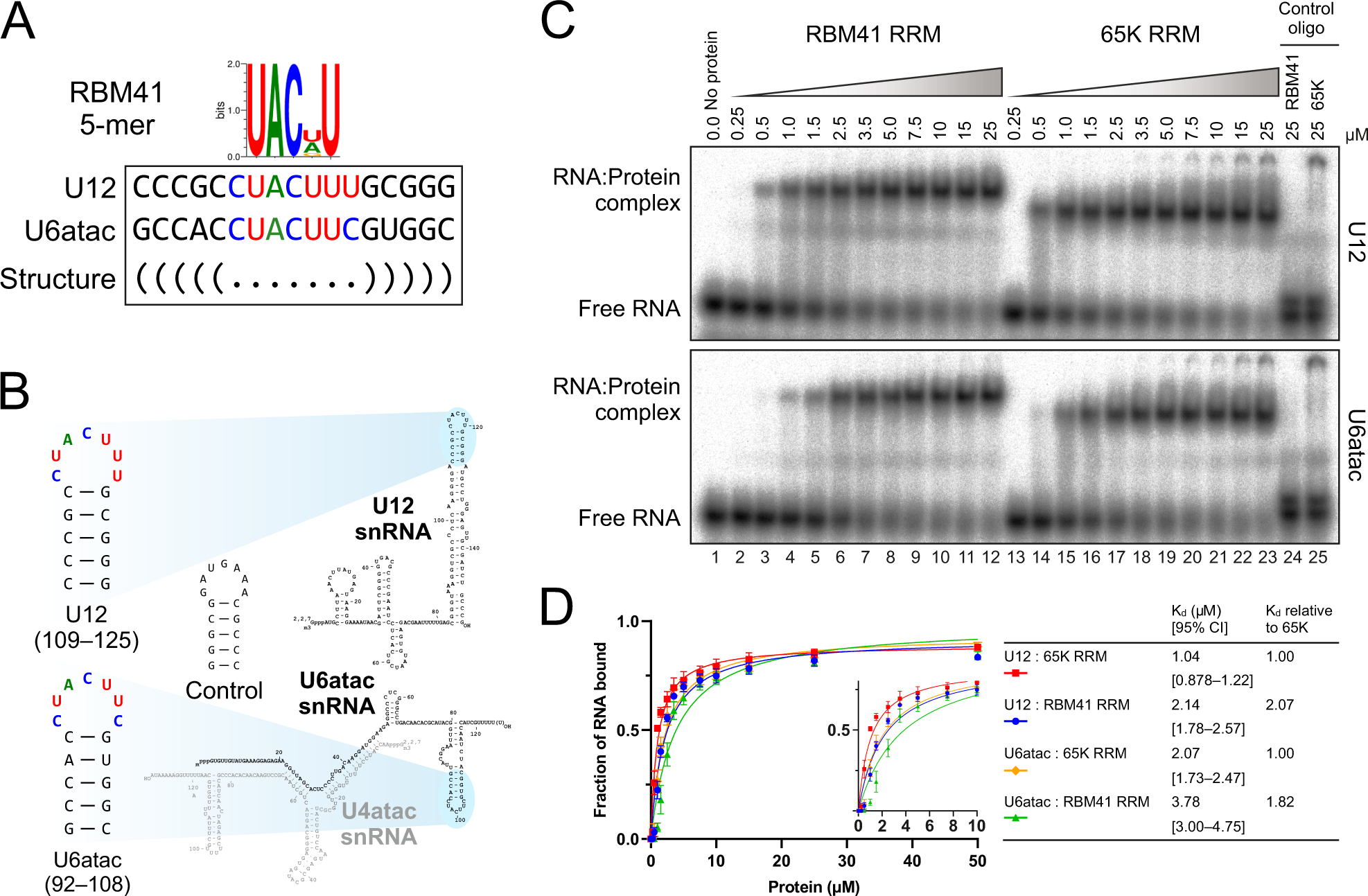
RBM41 interacts with the U12 and U6atac snRNAs *in vitro*. (A) Consensus RNA motifs bound by RBM41 *in vitro* and matching sequences in the U12 and U6atac snRNAs. The consensus motif determined by Ray et al. (Ray *et al*., 2013) using the RNAcompete method is shown. (B) RNA hairpins used in EMSA experiments and their location in the U12 and U6atac snRNAs. (C) EMSA analysis of RBM41 and U11/U12-65K RRM binding to U12 (top panel) and U6atac snRNA (bottom panel) hairpins. EMSA was carried out using recombinant RBM41 RRM (residues 267–413) or 65K C-terminal RRM (residues 380–517) and ^32^P-labeled U12, U6atac or negative control RNA hairpins shown in panel B. (D) Binding curves and dissociation constants for the interaction of RBM41 and 65K RRMs with U12 and U6atac hairpins. The inset shows a low protein concentration range (0– 10 µM) of the same binding curves.

Our EMSA analyses revealed that U11/U12-65K C-RRM and RBM41 RRM have similar overall binding characteristics, binding both to U12 and U6atac hairpins (Figure 2C, compare lanes1–12 and 13–23). In contrast, no complex formation was observed with either RRM when a control hairpin (complementary to the U12 hairpin) was used as a ligand (lanes 24 and 25). A further determination of the dissociation constants (K_d_) revealed that the RBM41 RRM has a low micromolar affinity to both U12 (K_d_=2.14 µM) and U6atac hairpins (K_d_=3.78 µM; Figure 2D). The 65K C-RRM showed approximately 2-fold higher affinity to both hairpins (U12: K_d_=1.04 µM; U6atac: K_d_=2.07 µM). Furthermore, both RRMs bound the U12 hairpin with ∼2-fold higher affinity compared to the U6atac hairpin (Figure 2D). Thus, both RBM41 and U11/U12-65K C-terminal RRMs show dual snRNA binding specificity *in vitro* with only slight differences in RNA affinity.

### RBM41 specifically associates with minor spliceosomal snRNPs

We next asked if RBM41 also associates with spliceosomal snRNAs *in vivo.* We transfected HEK293 cells with V5-tagged 65K or RBM41 followed by RNA immunoprecipitations (RIP) with anti-V5 antibody and northern blot analysis of minor and major spliceosomal snRNAs (Figure 3A). Consistent with its function in the U11/U12 di-snRNP, V5-65K strongly co-immunoprecipitated the U11 and U12 snRNAs and to a lesser extent U6atac and U4atac snRNAs (Figure 3A, lane 8). In contrast, V5-RBM41 co-immunoprecipitated the U12 snRNA, U4atac and U6atac snRNAs, but not the U11 snRNA (Figure 3A, lane 7). Similar results were obtained with an antibody against the endogenous RBM41 in HeLa nuclear extract (Figure 3B, lanes 1–4) or HEK293 total cell lysate (lanes 5–8), showing that these interactions are not artifacts resulting from RBM41 overexpression. U1, U2, U4, U5 and U6 snRNAs were variably detected slightly above control IP levels with transiently overexpressed V5-RBM41 (Figure 3A,D) and endogenous RBM41 (Figure 3B), likely due to non-specific association of RBM41 with these highly abundant snRNPs (see below).

**Figure 3.**
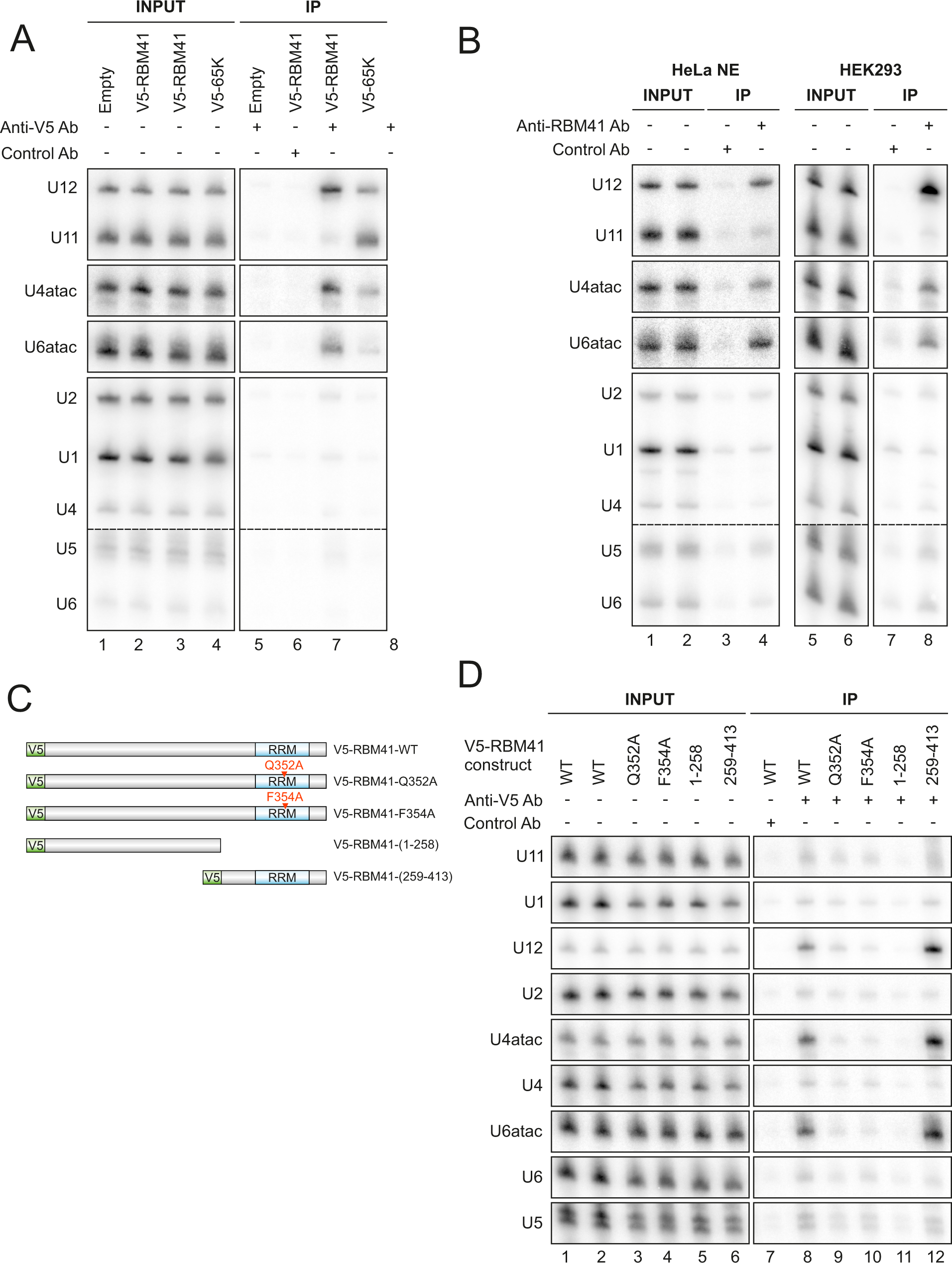
RBM41 specifically associates with minor spliceosomal snRNPs. (A) RNA immunoprecipitation with V5-tagged RBM41 and 65K. V5-RBM41 or V5-65K expression vector or empty vector were transfected into HEK293 cells. 24 h later, RNA immunoprecipitation with anti-V5 antibody or control antibody was carried out in native conditions and co-immunoprecipitated RNA analyzed by northern blot using the indicated probes. (B) RNA immunoprecipitation with endogenous RBM41. RIP was carried out in native conditions in either HeLa nuclear extract (left) or HEK293 total lysate (right) using an antibody against endogenous RBM41 or control antibody. (C) V5-RBM41 constructs used for RNA immunoprecipitation in panel D. (D) Effect of truncations and RRM mutations on the snRNP association of RBM41.V5-tagged RBM41 constructs shown in *C* were transfected into HEK293 cells and RNA immunoprecipitation carried out using anti-V5 or control antibody.

Next, we tested the role of the RBM41 RRM and the large N-terminal region lacking any identifiable domains in the snRNP association of RBM41. We carried out RIP experiments using wild-type V5-RBM41, V5-RBM41 lacking the RRM domain (V5-RBM41(1–258)) or the entire N-terminal region (V5-RBM41(259–413)), as well as V5-RBM41 constructs with alanine substitutions of Gln352 or Phe354 of the conserved YQF triad in the RRM (V5-RBM41-Q352A and V5-RBM41-F354A) (Figure 3C). The Q352A and F354A mutations and deletion of the RRM led to dramatic loss of the U12, U4atac and U6atac interactions (Figure 3D, lanes 8–11), while the V5-RBM41(259–413) construct still showed robust co-immunoprecipitation of all three snRNAs (lane 12). Notably, while major spliceosome-specific snRNAs (U1, U2, U4, U6) and the shared U5 snRNA were also detected above control IP levels in the V5-RBM41 anti-V5 IP (lane 8), these were unaffected by the RRM mutations or the truncations, indicating that these IP signals represent nonspecific background rather than specific interactions. Taken together, our RIP experiments show that RBM41 specifically associates with minor spliceosomal snRNPs in the cell and suggest that the snRNP association of RBM41 is primarily mediated through the RRM binding with its target snRNAs.

### RBM41 and U11/U12-65K partition into distinct snRNP complexes

While our *in vitro* binding experiments demonstrated similar RNA-binding properties for RBM41 and U11/U12-65K, the distinct snRNA IP profiles and the lack of sequence similarity outside the RRM suggested functional divergence of the two proteins within the minor spliceosome. As a complementary method to study snRNP complex association of RBM41 and U11/U12-65K, we carried out glycerol gradient fractionation of HeLa nuclear extract and analyzed the sedimentation behavior of the two proteins by western blot and spliceosomal snRNAs by northern blot (Figure 4A). Consistent with our RIP experiments, RBM41 and U11/U12-65K were largely found in different fractions, indicating association with different molecular complexes. RBM41 peak co-migrated with the U12 mono-snRNP (Figure 4A, fractions 8–10), whereas the U11/U12-65K peak co-migrated with the U11/U12 di-snRNP (fractions 12–14). Though the U4atac or U6atac snRNAs did not show clear co-migration with RBM41, a minor fraction of these snRNAs was present in RBM41 peak gradient fractions. Taken together with our RIP data, while the two proteins share similar RNA binding specificity *in vitro*, in the cell they partition into distinct snRNP complexes.

**Figure 4.**
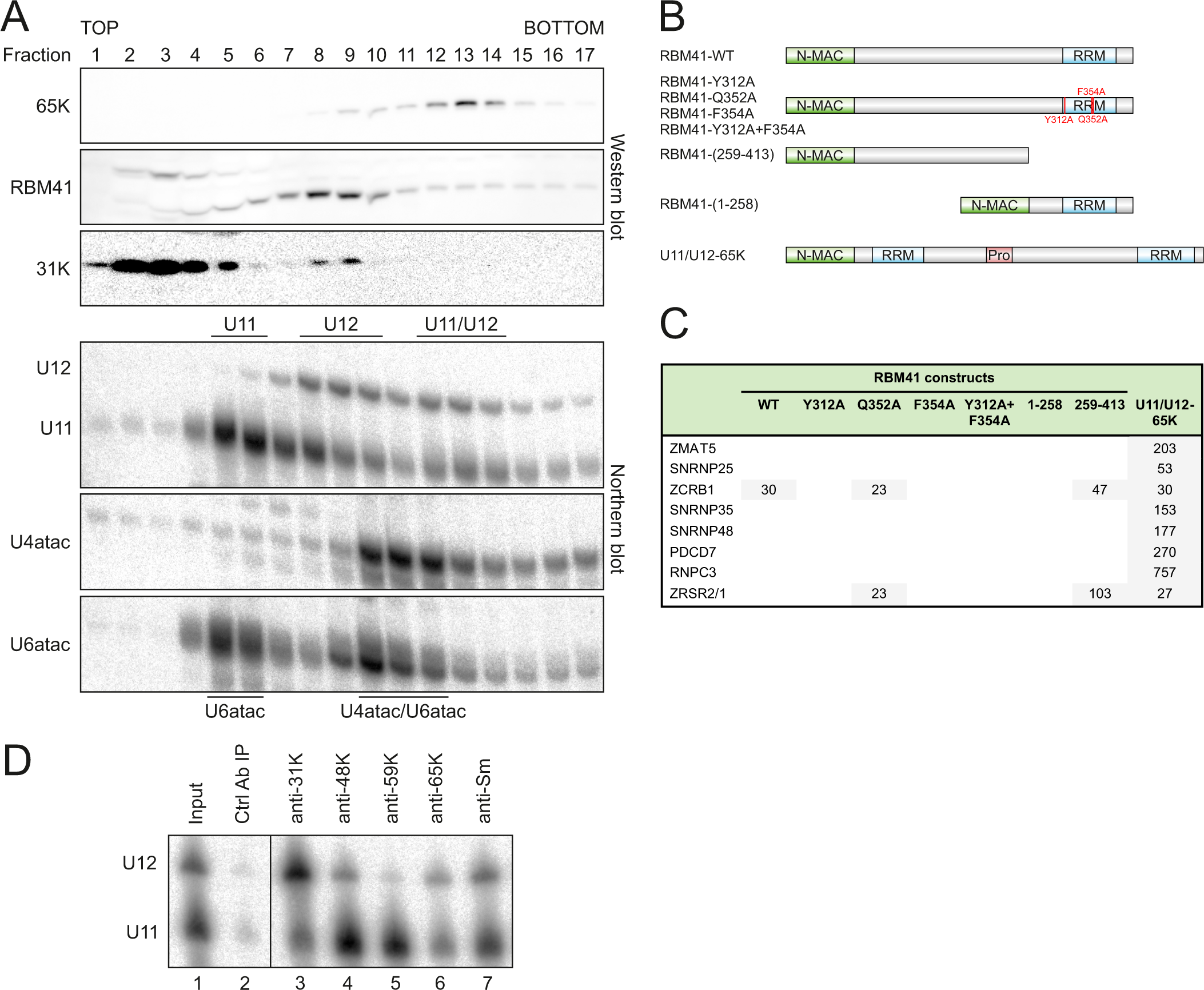
RBM41 and U11/U12-65K partition into distinct snRNP complexes. (A) Glycerol gradient analysis of RBM41 and U11/U12-65K in HeLa nuclear extract. Nuclear extract was loaded on top of a 10–30% glycerol gradient. After ultracentrifugation, the gradient was fractionated, protein and RNA isolated and analyzed by western and northern blot using the antibodies and probes indicated on the left. Location of the U11, U12 and U6atac mono-snRNPs, U11/U12 di-snRNP and U4atac/U6atac di-snRNP are inferred based on the snRNA profiles. (B) Domain structures of MAC-tagged RBM41 and 65K constructs used for BioID. N-terminal MAC tag is not drawn to scale. (C) Spectral count fold changes for U11/U12 di-snRNP proteins in BioID datasets. (D) Immunoprecipitation of U11 and U12 snRNAs by anti-31K, anti-48K, anti-59K, anti-65K and anti-Sm antibodies in HEK293 total lysate followed by Norther blot detection of the U11 and U12 snRNAs.

To define the composition and interactions of the RBM41 in the U12 mono-snRNP related to the 65K in U11/U12 di-snRNP, we used proximity-based labeling (BioID) to map the proximity interactors of the two proteins. A panel of RBM41 constructs and a U11/U12-65K construct (Figure 4B), each carrying an N-terminal MAC-tag (consisting of BirA* biotin ligase, hemagglutinin (HA) and StrepIII tags), were integrated into the Flp-In™ T-REx™ 293 cell line, enabling both inducible transgene control (tetracycline) and inducible biotinylation of proteins coming into proximity with the bait protein. Wild-type U11/U12-65K, wild-type RBM41, four RNA binding deficient RBM41 mutants (Y312A, Q352A, F354A, Y312A+F354A) and two RBM41 truncation constructs (1–258 and 259–413) were analyzed (Figure 4B). Western blot analysis with anti-HA antibody confirmed correct expression of MAC-tagged bait proteins (Supplementary Figure 3A), while biotinylation was visualized with avidin-HRP (Supplementary Figure 3B). Immunofluorescence with anti-HA was used to detect localization of the MAC-tagged constructs (Supplementary Figure 4). MAC-RBM41-WT and MAC-U11/U12-65K constructs showed a predominantly nuclear localization, with MAC-RBM41-WT showing a more prominent cytoplasmic subpopulation. In contrast, MAC-RBM41-(1-258) was almost uniformly distributed between cytoplasm and nucleus, suggesting that deletion of the C-terminal RRM impairs nuclear import of RBM41.

MAC-tagged protein expression was induced with tetracycline in the presence of biotin for 24 hours before harvesting of cells. Three independent replicates were analyzed for each cell line using an established BioID pipeline that has been described in detail (Liu *et al*., 2018; Liu *et al*., 2020). Consistent with its role in U11/U12-di-snRNP, MAC-U11/U12-65K interacted with all 8 minor spliceosome-specific proteins of the U11/U12 di-snRNP complex (Figure 4C, Supplementary Table 4), with its top interactors being U11-59K (PDCD7), U11/U12-20K (ZMAT5), U11-48K (SNRNP48) and U11-35K (SNRNP35). In contrast, MAC-RBM41-WT only interacted with one of the U11/U12 di-snRNP proteins, U11/U12-31K (ZCRB1) (Figure 4C, Supplementary Table 4). This interaction was dependent on a functional RBM41–U12 snRNA interaction as the construct encoding the RRM-only version of MAC-RBM41(259– 413) was still able to support the interaction with U11/U12-31K while the RRM mutants and the RRM deletion construct (MAC-RBM41-(1–258)) did not. Additionally, the loss of the RBM41 N-terminal domain resulted in a strong interaction with ZRSR1/2, which functions in 3’ss recognition of U12-type introns.

A further support for the direct or indirect interaction between RBM41 and U11/U12-31K within the U12 mono-snRNP is obtained from glycerol gradient centrifugation which shows that RBM41 and and U11/U12-31K proteins cosediment in the same U12 mono-snRNP fractions (Figure 4A, lanes 8-9). Similarly, co-IP experiments show that the endogenous U11/U12-31K is preferentially associated with U12 snRNA, unlike the U11/U12-65K which shows an even co-IP efficiency for both U11 and U12 snRNAs (Figure 4D, cf. lanes 3 and 6). Association of the U11/U12-31K with the mono-snRNP is consistent with direct recognition of 2’*-O-* methylated A8 residue of the U12 snRNA (Li *et al*., 2023). Conversely, the inverted co-IP pattern with U11-48K and U11-59K is consistent with their role as components of both U11 mono-snRNP and the U11/U12 di-snRNP (Fig 4D, lanes 4 and 5). Together, our RIP, glycerol gradient and BioID data show that RBM41 and U11/U12-65K partition into distinct snRNP complexes and are likely to play distinct roles within the minor spliceosome.

### RBM41 interacts with DHX8 and localizes to Cajal bodies

One of the top BioID proximity-labeling hits of MAC-RBM41-WT was the DEAH-box RNA helicase DHX8 (Figure 5, Supplementary Table 4). The interaction with DHX8 was dependent on the highly conserved N-terminal region of RBM41, while the RRM mutations and deletion of the RRM had no effect. In contrast, no interaction of U11/U12-65K with DHX8 was detected in BioID analysis. The DHX8:RBM41 interaction was also reported in a recently published reciprocal DHX8 BioID dataset (Go *et al*., 2021).

**Figure 5.**
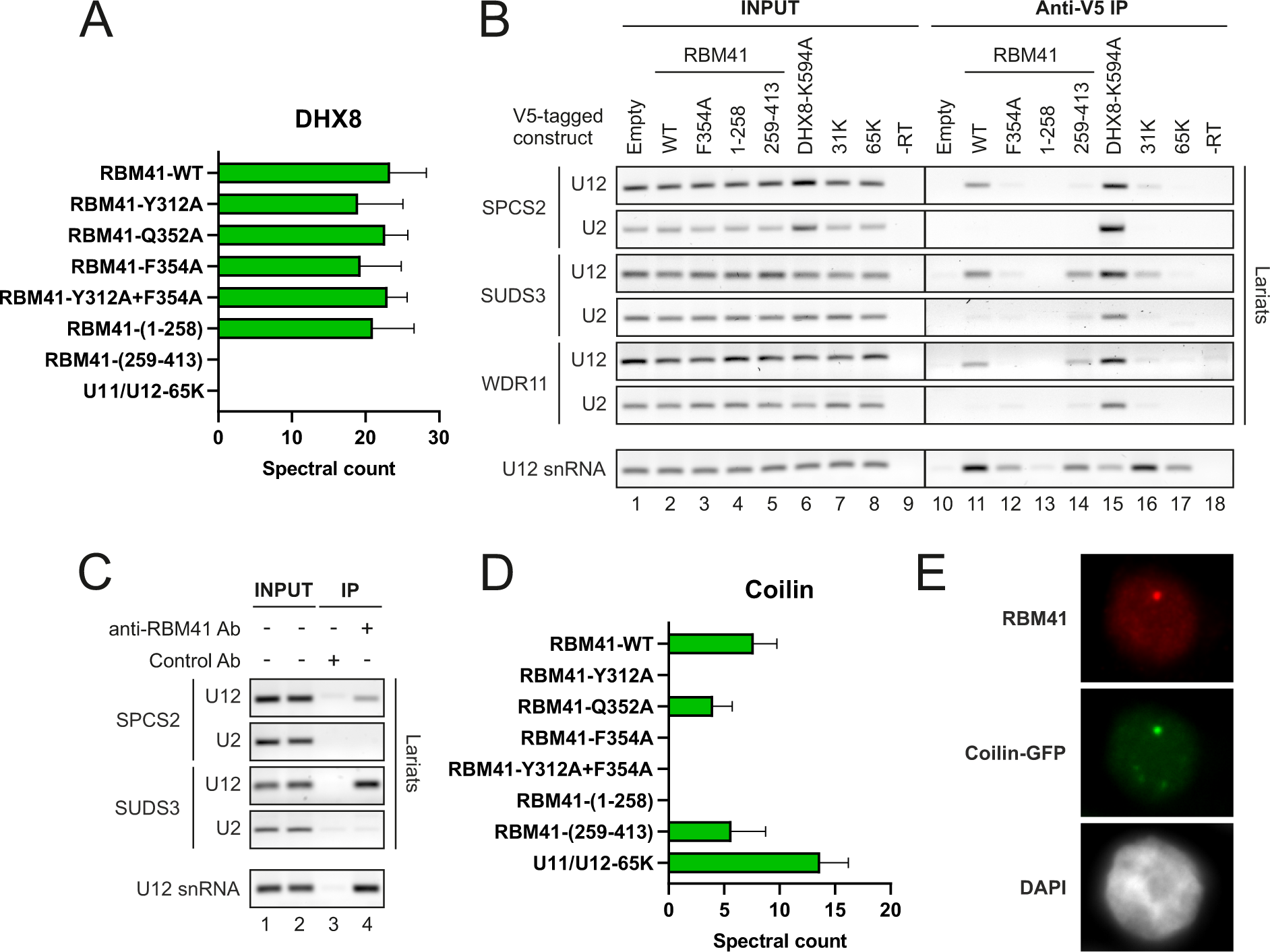
RBM41 interacts with DHX8 and localizes to Cajal bodies. (A) Spectral counts for DHX8 in RBM41 and U11/U12-65K BioID datasets. (B) RNA immunoprecipitation with exogenously expressed V5-tagged proteins followed by RT-PCR. The indicated pCI-neo constructs for expressing V5-tagged proteins or empty pCI-neo vectors were transfected into HEK293 cells. 24 h later, RIP was carried out using anti-V5 antibody and RNA extracted from the beads analyzed by RT-PCR. Amplification across the branch junction was used to detect U2- and U12-type intron lariats and lariat intermediates from the following introns: SPCS2 introns 3–4 (U12) and 2–3 (U2), SUDS3 introns 7–8 (U12) and 9–10 (U2), WDR11 introns 28–29 (U12) and 27-28 (U2). (C) RNA immunoprecipitation with endogenous RBM41 in HEK293 cells followed by RT-PCR. (D) Spectral counts for coilin (COIL) in RBM41 and U11/U12-65K BioID datasets. (E) Anti-RBM41 immunofluorescence in HEK293 cells transfected with a vector for expressing coilin-GFP.

DHX8 has not been shown to function in the minor spliceosome. In the major spliceosome, it is recruited before exon ligation, during the transition from C to C* complex (Zhan *et al*., 2018), and drives the P-to-ILS1 transition presumably by pulling on the ligated exon, leading to its release and dissociation of at least nine proteins (Zhang *et al*., 2019). We thus hypothesized that RBM41 could have a similar function in the late stages of minor splicing, possibly in the post-catalytic complexes. To test association of RBM41 and DHX8 with late-stage minor spliceosomes, we carried out a RIP experiment in HEK293 cells transfected with V5-RBM41, V5-RBM41 mutant and truncation constructs (F354A, 1-258 and 259-413), or V5-tagged DHX8 carrying a helicase mutation (K594A) that stalls splicing in the P complex stage (Fica *et al*., 2019; Figure 5B; Ohno *et al*., 1996). V5-65K and V5-31K were analyzed as controls for U11/U12 proteins. As DHX8 associates with post-branching spliceosomes, we used RT-PCR across branch sites (Lorsch *et al*., 1995; Suzuki *et al*., 2006) to detect U2-type and U12-type lariat intermediates and excised intron lariats in the immunoprecipitation. V5-DHX8-K594A co-immunoprecipitated both U2-type and U12-type lariats from the same genes (Figure 5B, lane 15), suggesting that DHX8 functions in both spliceosomes. In contrast, RBM41 preferentially co-immunoprecipitated U12-type over U2-type intron lariats (lane 11). Similar results were obtained when RIP was carried out using an antibody against the endogenous RBM41 protein (Figure 5C). While U11/U12-65K did not co-IP any lariats (lane 17), the U11/U12-31K shows variable and low levels of lariat co-IP (lane 16), suggesting that RBM41 BioID hits for DHX8 and U11/U12-31K may be originating from two separate complexes.

Another major interactor of MAC-RBM41-WT in our BioID data was coilin (Figure 5D, Supplementary Table 4), a key scaffolding protein and widely used marker for Cajal bodies. Consistently, we found that endogenous RBM41 localizes to Cajal bodies, as shown by colocalization with coilin-GFP in HEK293 cells (Figure 5E). While U11/U12-65K also interacted with coilin, the nuclear bodies labeled by the anti-RBM41 antibody did not colocalize with endogenous U11/U12-65K (Supplementary Figure 4B). The coilin:RBM41 interaction was dependent on U12 snRNA binding, as mutating or deleting the RBM41 RRM reduced or completely eliminated the interaction, while MAC-RBM41-(259-413) was still able to interact with coilin (Figure 5D). Similarly, anti-HA immunofluorescence staining in BioID cell lines detected nuclear bodies in cells expressing MAC-RBM41-WT, but not in cells expressing any of the RBM41 RRM mutants or MAC-RBM41-(1-258) (Supplementary Figure 4A). This suggests that RBM41 localizes to Cajal bodies in an RNA binding-dependent manner.

### RBM41 is not essential for cell viability but affects the splicing of a subset of U12-type introns

To assess the effect of RBM41 loss on the splicing of the U12-type introns, we generated several independent RBM41 full knockout lines with HEK293 cells using CRISPR-Cas9 editing. The loss of functional RBM41 loci in each of the three X chromosomes in HEK293 cells was confirmed by Sanger sequencing and western blot analyses (Fig 6A; Supplementary Figure 5). The knockout cells were not only viable, but the loss of RBM41 did not lead to any noticeable growth phenotypes. This is consistent with the earlier investigations on human essentialomes that consistently indicated that RBM41 locus is not essential for the cell viability (Blomen *et al*., 2015; Yilmaz *et al*., 2018).

**Figure 6.**
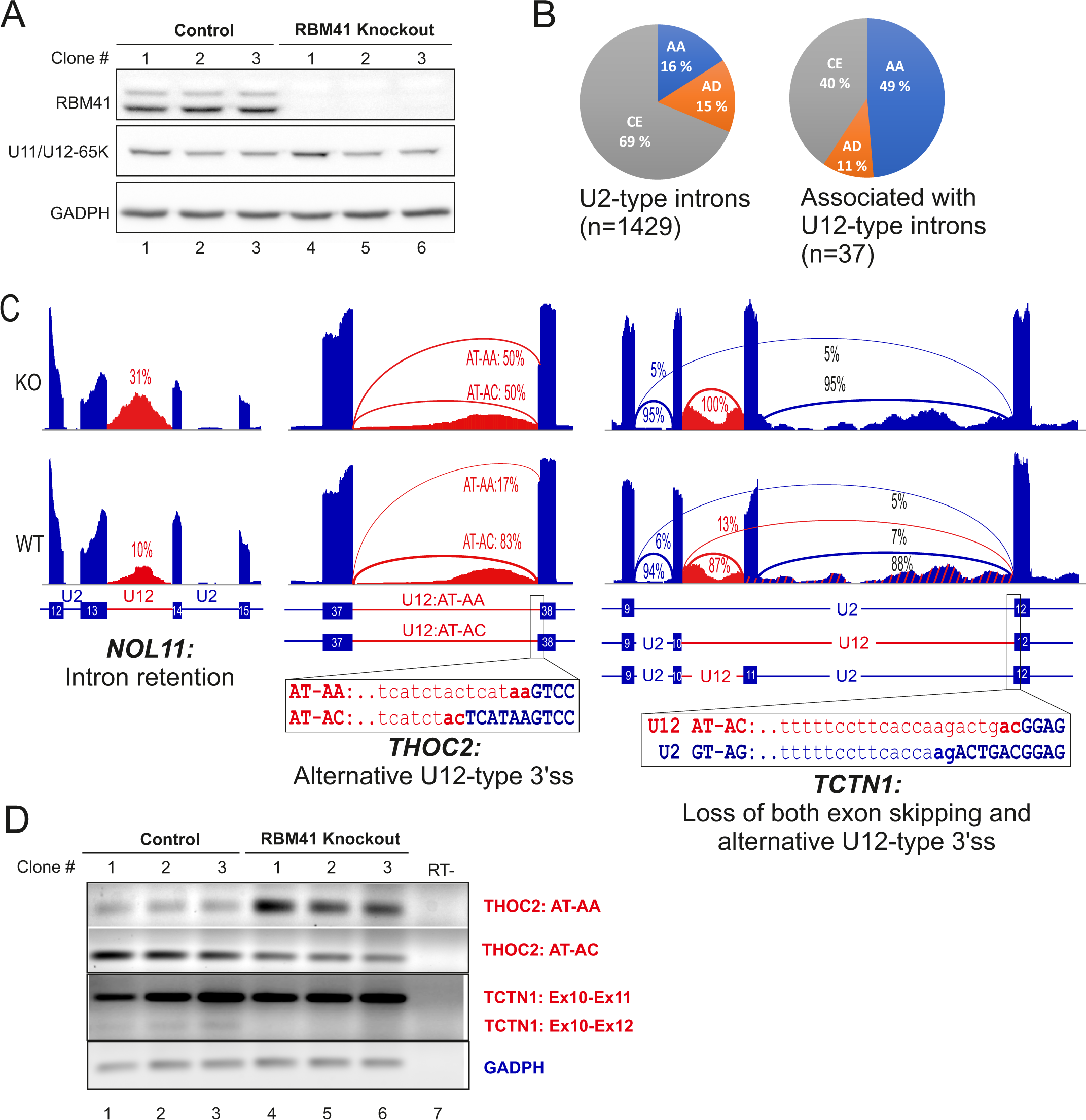
RBM41 knockout influences the splicing of U12-type introns. (A) Western blot analysis of RBM41 knockout and matching control cell lines used in the RNASeq analysis. (B) Comparison of the statistically significant (Whippet Probability > 0.9) alternative splicing events in the genes containing only U2-type introns and events either within or near proximity (immediate up- or downstream exons and introns) of the U12-type introns. AA - alternative acceptor, AD - alternative donor, CE - Cassette exon. (C) Representative sashimi plots showing Intron retention (*NOL11*), Alternative U12-type 3’ss choice (*THOC2*) and loss of both exon skipping and alternative U12-type 3’ss usage in RBM41 knockout cells (*TCTN1*). (D) validation of the *THOC2* and *TCTN1* alternative splicing changes using a set of three independent RBM41 knockout cell lines and their matching controls.

To analyze the effects of RBM41 knockout on splicing we carried out RNAseq analysis of three independent knockout cell lines and matching unedited lines. Subsequent bioinformatics analysis concentrated on U12-type intron retention and alternative/cryptic splice site activation with the U12-type intron containing genes. Intron retention analysis revealed weak splicing defects in the knockout cell lines for a small subset of genes (14 genes), such as the *NOL11* (Fig 6C; Table S6), but also identified 15 genes that instead showed the opposite, that is, a reduction in the read levels mapping to the U12-type introns (Table S6). Additionally, we investigated alternative splice donor (AD), splice acceptor (AA) and cassette exon (CE) usage. We further focused on the events within the U12-type introns and the surrounding exons and introns as these are the potential direct targets of RBM41 knockdown. We detected a total of 37 statistically significant alternative splicing events in 26 genes, as several genes showed multiple AS events being activated as a result of RBM41 knockout (Table S7). Of these, the most notable were the ∼3-fold enrichment in alternative 3’ss (AA) usage (Figs. 6B-D; p=3.4−10^−6^, hypergeometric test) and the 1.7-fold reduction in cassette exon (CE) events (Figs. 6B-D; p=1.2−10^−4^, hypergeometric test) when compared to the alternative splicing events detected in genes containing only major introns. Notably, most of the identified alternative 3’ss events (13/18) affected the U12-type intron 3’ss choice (Table S6), suggesting that the loss of RBM41 has a weak, but nevertheless statistically significant effect on the splicing of a subset of U12-type introns.

## DISCUSSION

In this work, we have expanded the repertoire of unique protein components specific to minor spliceosome by providing evidence that RBM41 functions in post-splicing steps of the minor spliceosome assembly/disassembly cycle. RBM41 shows a similar phylogenetic co-evolution pattern as several other minor spliceosome components (Figure 1E) and it has earlier been annotated as a paralog of U11/U12-65K protein, due to the highly similar C-terminal RRMs found in the two proteins. Here, we show that the C-terminal RRM of RBM41 binds to the 3’-terminal stem-loops of U12 and U6atac snRNAs both *in vitro* and *in vivo*. Compared to the U11/U12-65K C-terminal RRM, RBM41 has approximately 2x lower affinity to its RNA ligands. We further show that unlike U11/U12-65K, which is a component of the U11/U12 di-snRNP, RBM41 associates with a distinct U12 mono-snRNP. Both U12 mono-snRNP and U11/U12 di-snRNP complexes have been described previously (Montzka *et al*., 1988; Wassarman *et al*., 1992; Will *et al*., 2004), but the function or composition of the U12 mono-snRNP has not been studied further. Here, our ultracentrifugation and BioID analysis provides evidence that the U12 mono-snRNP is a distinct functional complex in the minor spliceosome and contains, in addition to RBM41, the U11/U12-31K (ZCRB1) protein as the specific protein components. Additionally, we show that RBM41 associates specifically with excised U12-type intron lariats and uses its unique N-terminal domain to interact with the DHX8 helicase, and likely cycles through the Cajal bodies. Together, our data suggests that the two paralogous proteins have distinct functions in U12-type intron splicing with U11/U12-65K functioning in the early steps of U12-type intron recognition and RBM41 in the post-splicing steps and during minor spliceosome disassembly.

Our results highlight the role of the 3’ stem-loop of U12 snRNA in the minor spliceosome assembly-disassembly cycle. The significance of the 5’ end of the U12 snRNA has long been recognized due to its function in the BPS recognition and the interactions with the U6atac snRNA in the catalytic core of the minor spliceosome (Turunen *et al*., 2013). In contrast, the 3’-terminal stem-loop of the U12 snRNA has appeared as a static binding site for the U11/U12-65K protein, necessary for the formation of the U11/U12 di-snRNP. Our identification of RBM41 binding to the 3’-terminal stem-loop during minor spliceosome disassembly suggests more dynamic recognition events where the 3’-terminal stem-loop serves as a platform for the two paralogous proteins which guide the U12 snRNA though the minor spliceosome assembly and disassembly cycle. The previously characterized steps include the recognition of the 3’-terminal stem-loop of the U12 snRNA by the U11/U12-65K protein, which uses its N-terminus to interact with the U11-59K protein (Benecke *et al*., 2005) to form the U11/U12 di-snRNP, which in turn is necessary for the U12-type intron recognition (Frilander *et al*., 1999). Furthermore, during the formation of the catalytically active spliceosome (B^act^ complex) the U11/U12-65K protein remain attached to the B^act^ complex (presumably to the 3’-terminal stem-loop), while U11 snRNP and all the other specific protein components of the di-snRNP are released from the activated spliceosome (Bai *et al*., 2021). Our data indicate that later in the splicing process there is an exchange in the 3’-terminal stem-loop binding partner from U11/U12-65K to RBM41 which can be detected in post-splicing complexes containing excised minor intron lariats, and which is also in close proximity with the DHX8/hPrp22 helicase. Together these results suggest that RBM41 is present in the minor spliceosome post catalytic (P) and intron lariat spliceosome (ILS) complexes. Furthermore, proximity labeling of Cajal body marker coilin (Figure 5D; Go *et al*., 2021) and co-localization of RBM41 and coilin in Cajal bodies (Figure 5E, Supplementary Figure 4) suggests that RBM41 remains bound to the U12 snRNA during the entire spliceosome disassembly process and follows the snRNA to the Cajal body. This is the likely location for a new round of assembly of the U11/U12 di-snRNP complex. Proximity interaction between RBM41 and U11/U12-31K (Figure 4C) and the association of U11/U12-31K with the U12 mono-snRNP (Figures 4A, D) suggest that the new round of U11/U12 snRNP assembly is initiated by the binding of U11/U12-31K to the 2’*-O-*methylated A8 position of U12 snRNA (Li *et al*., 2023), followed by the exchange from RBM41 to U11/U12-65K.

RBM41 and U11/U12-65K proteins interact with both U12 and U6atac 3’ terminal stem-loops *in vitro* and *in vivo*. However, the U12 interactions appear more significant, given the sensitivity of U12-type intron splicing to mutations in the U12 single-stranded loop and the insensitivity to U6atac loop mutations (Sikand *et al*., 2011; Singh *et al*., 2016). Furthermore, an 84C>U mutation that compromises the U12 3’-terminal stem-loop integrity leads to early onset cerebellar ataxia due to overtrimming of the 3’-terminal stem-loop which removes the binding site of the U11/U12-65K and RBM41 proteins (Elsaid *et al*., 2017; Norppa *et al*., 2021). Similarly, the U11/U12-65K P474T mutation associated with isolated growth hormone deficiency has been shown to reduce the U11/U12 di-snRNP levels due to a folding defect of the U11/U12-65K C-RRM, which reduces its affinity to the 3’-terminal stem-loop (Argente *et al*., 2014; Norppa *et al*., 2018). However, in that case the potential additional effects on U6atac binding *in vivo* or on the recycling of the U12 snRNA have not been ruled out.

Our data portrays a somewhat conflicting view on the significance of the RBM41 and the need for a specific protein factor(s) for the minor spliceosome disassembly process. The strong sequence conservation observed with the domains of RBM41 that interact either with the U12 snRNA or DHX8 (Figure S2) suggests a strong selection pressure at the organismal level to maintain these interactions. This is reflected in phylogenetic co-occurrence in multiple evolutionary lineages (Figure 1E), though we note that the locus encoding RBM41 is more frequently absent in multiple evolutionary lineages than several other components of the minor spliceosome. On the other hand, both our knockout data (Figure 6A) and data from essentialomes (Blomen *et al*., 2015; Yilmaz *et al*., 2018) indicate that RBM41 is dispensable at least at the cellular level. Based on the weak, but yet statistically significant effects of RBM41 knockout in human HEK293 cells specifically on 3’ss selection of U12-type introns (Figure 6B-D), we hypothesize that while RBM41 is dispensable at the cellular level, it may nevertheless be able to exert a weak kinetic effect on splicing in addition to later participating in the disassembly process. The effect on 3’ss choice is similar to that observed after major spliceosome catalytic step II factor knockdowns, which similarly influence the 3’ss choice, particularly with NAGNAG introns (Dybkov *et al*., 2023), further suggesting that the exchange from the U11/U12-65K to RBM41 may take place prior to step II. However, given that our BioID analysis did not provide supporting evidence for this possibility, it is also possible that the exchange from U11/U12-65K to RBM41 takes place at a later step and the effects on minor intron splicing are secondary effects of downstream processes being disturbed. Finally, while RBM41 protein is not absolutely needed for cell viability or splicing in the highly proliferative cell types used in essentialome and our knockout studies, it may provide selective advantage in specific cell types or in the context of whole organisms to account for the observed evolutionary conservation.

RBM41 may also play a role in substituting structures or interactions that are present in the major but not in the minor spliceosome. Specifically, the human minor B^act^ complex lacks several key protein components that are present in the major B^act^ complex. These include NTC complex proteins (PRPF19, SPF27 and SYF1), NTR complex proteins (BUD31 and RBM22), SF3a complex, and phosphoprotein isomerases (PPIL1 and CypE). Conversely, the minor B^act^ complex contains four unique proteins RBM48, ARMC7, SCNM1 and CRIPT that are not present in the major B^act^ complex (Bai *et al*., 2019; Bai *et al*., 2021; Siebert *et al*., 2022). At least a subset of these differences in protein composition can be explained by divergent snRNA components between the spliceosomes which change the RNA elements available for the binding of protein factors. For example, U6atac snRNA lacks the 5’ stem-loop structure which is present in U6 snRNA and recognized by the BUD31 and RBM22 proteins in major B^act^ complex. Instead, in minor B^act^ these proteins are replaced by RBM48 and ARMC7 proteins binding to the γ-monomethyl cap of U6atac snRNA (Bai *et al*., 2021). Similarly, as the U11/U12 di-snRNP lacks the SF3a complex (Will *et al*., 1999) that is present both in the 17S U2 snRNP and the major B^act^ complex (Bessonov *et al*., 2010; Schmidt *et al*., 2014), the functions of SF3a in minor B^act^ has been taken over by the SCNM1, a unique protein component of the minor spliceosome (Bai *et al*., 2021). In this respect the absence of CypE and SYF1 from the minor B^act^ (Bai *et al*., 2021) is intriguing as those are loaded to major B^act^ and are later involved in post-spliceosomal complex transitions (Fica *et al*., 2019; Zhang *et al*., 2019). From the available limited structural information, we hypothesize that there may be similar differences in the post-spliceosomal complex composition between the two spliceosomes such that RBM41 may substitute some of the major spliceosome structures and /or interactions that are missing from the minor spliceosome. Testing of this hypothesis would require similar high-resolution structures of the post-splicing minor spliceosome complexes that are available for the major spliceosome.

Unique protein components in the minor spliceosome can also offer potential opportunities for differential regulation of the two spliceosomes. Given the function of RBM41 in the post-splicing complexes, the regulation of minor intron splicing via RBM41 abundance or activity is unlikely. However, earlier work has nevertheless provided evidence of translational regulation of RBM41 via differential 3’ UTR isoform usage (Sterne-Weiler *et al*., 2013), particularly during neuronal development (Yap *et al*., 2016). As the abundance of U11/U12-65K is also regulated by a feedback/cross-regulation pathway, particularly during neuronal differentiation (Verbeeren *et al*., 2010; Verbeeren *et al*., 2017), it is possible that the cellular levels of RBM41 protein are similarly linked to the abundance of minor spliceosome components or to global regulation of mRNA processing pathways.

## Acknowledgements

We thank the members of the Frilander lab for suggestions during the work and Marja-Leena Peltonen for technical assistance.

---

**Supplementary Figure 1.**
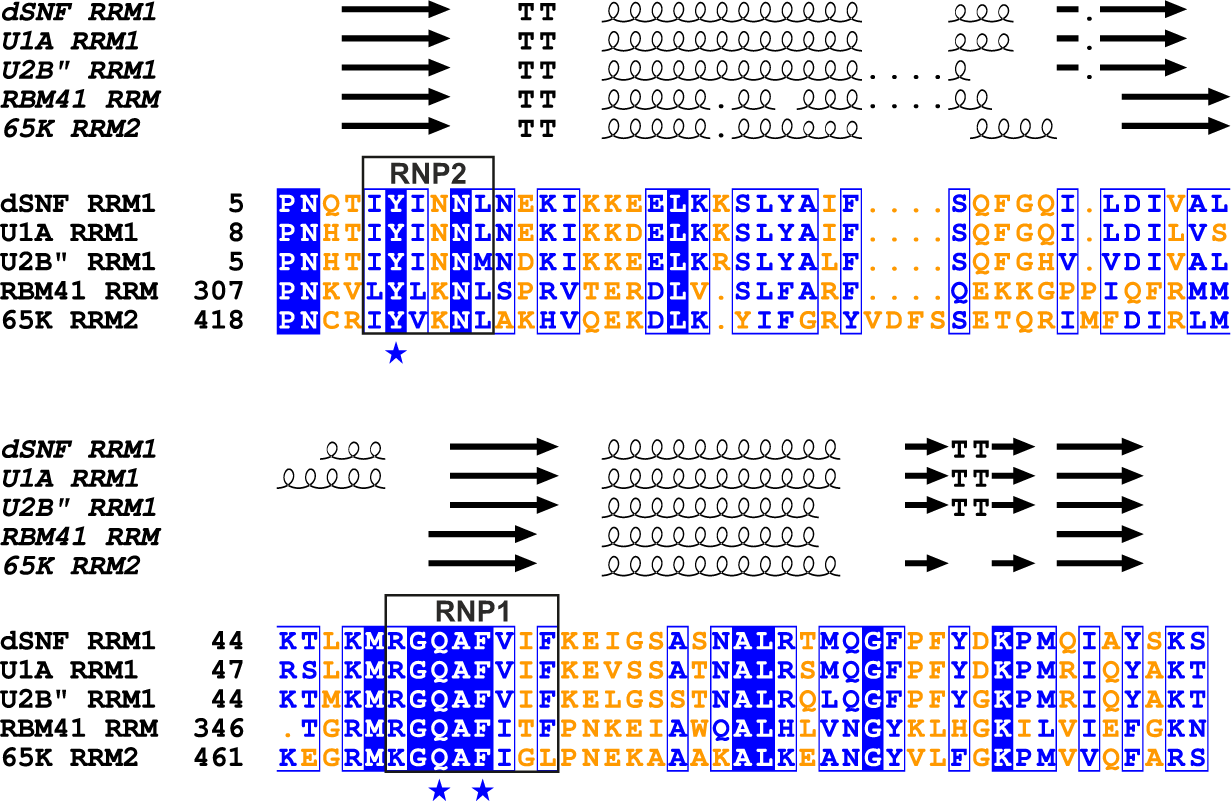
Multiple sequence alignment of human RBM41, U11/U12-65K, U1A, U2B″ and *Drosophila melanogaster* SNF (sans fille) RNA recognition motifs. Alignment was carried out using MAFFT and visualized using ESPript 3.0. Protein secondary structure information was extracted from the following PDB structures: 6F4I, 1NU4, 1A9N, 2CPX, 5OBN. The conserved YQF triad residues are indicated with blue stars.

**Supplementary Figure 2.**
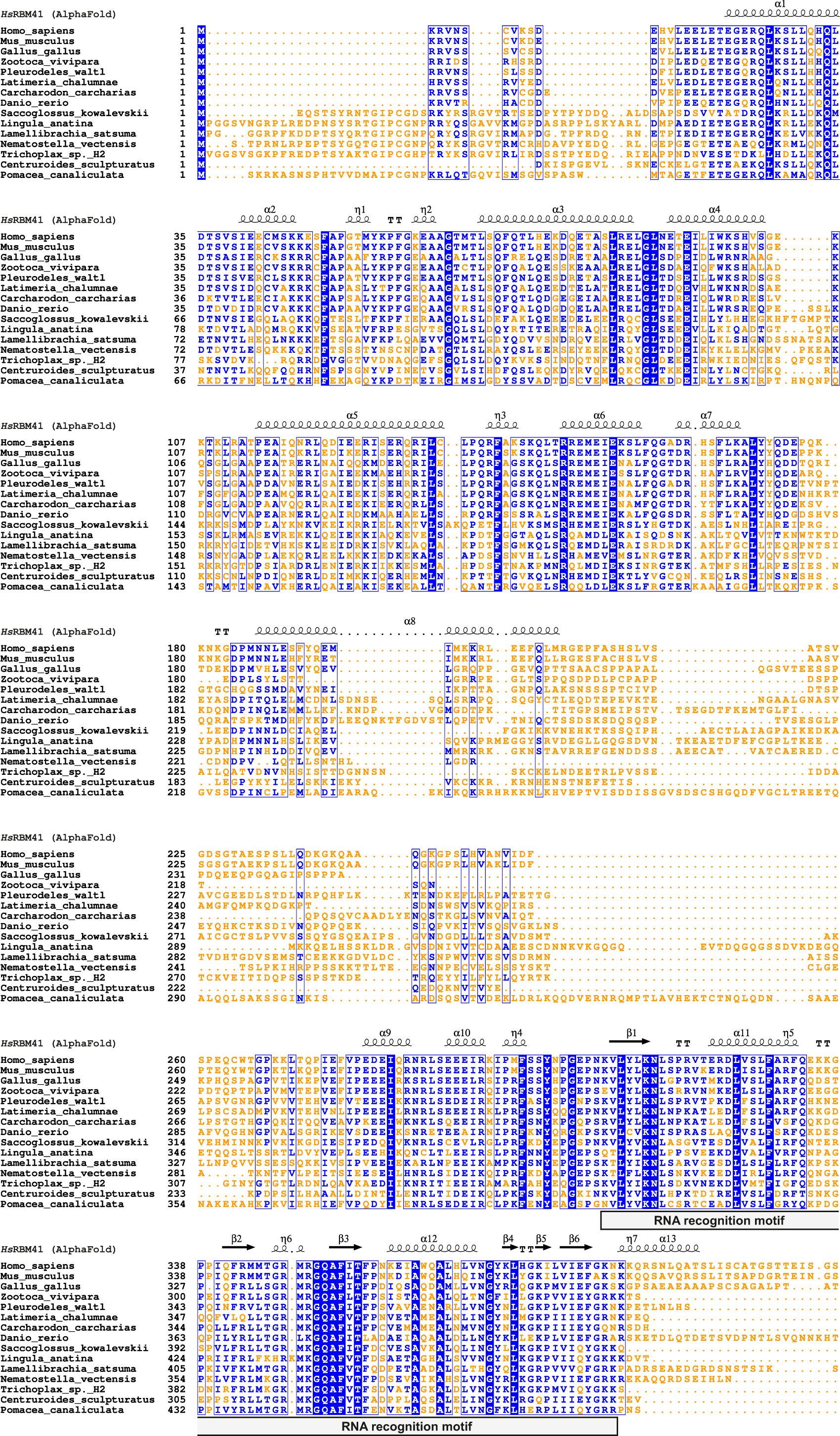
Multiple sequence alignment of RBM41 orthologs from 15 animal species. Alignment was carried out using MAFFT and visualized using ESPript 3.0. Secondary structures extracted from AlphaFold-predicted human RBM41 structure (Figure 1D) are shown. Sequences used for the alignment are listed in Supplementary Table 2.

**Supplementary Figure 3.**
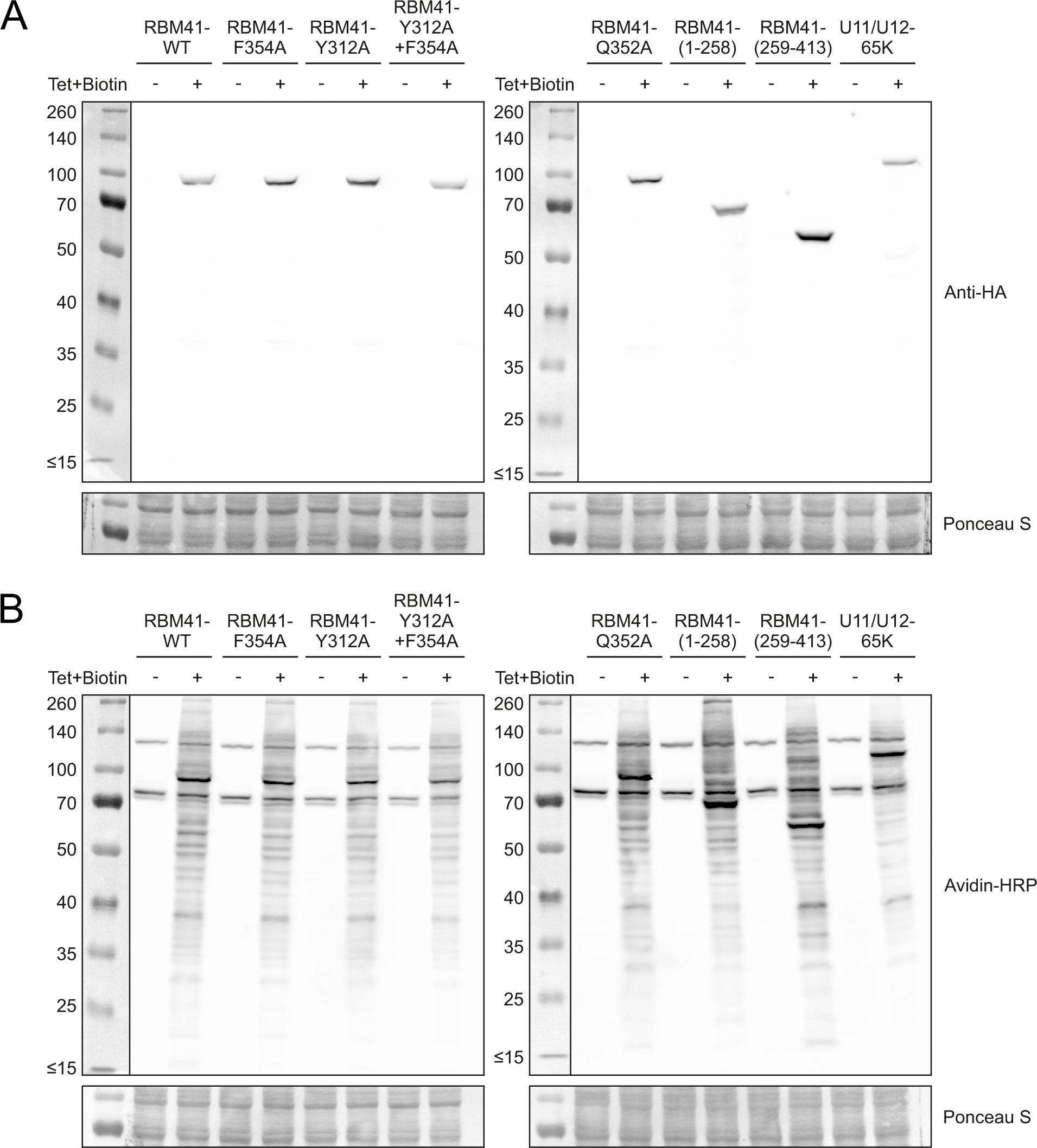
Validation of Flp-in 293 cell lines used in BioID experiments. MAC-tagged protein expression and biotinylation was induced by addition of tetracycline and biotin for 24 hours before harvesting of cells. **(A)** Induction of MAC-tagged proteins detected by western blot with anti-HA antibody. **(B)** Biotinylation detected by western blot with avidin-HRP.

**Supplementary Figure 4.**
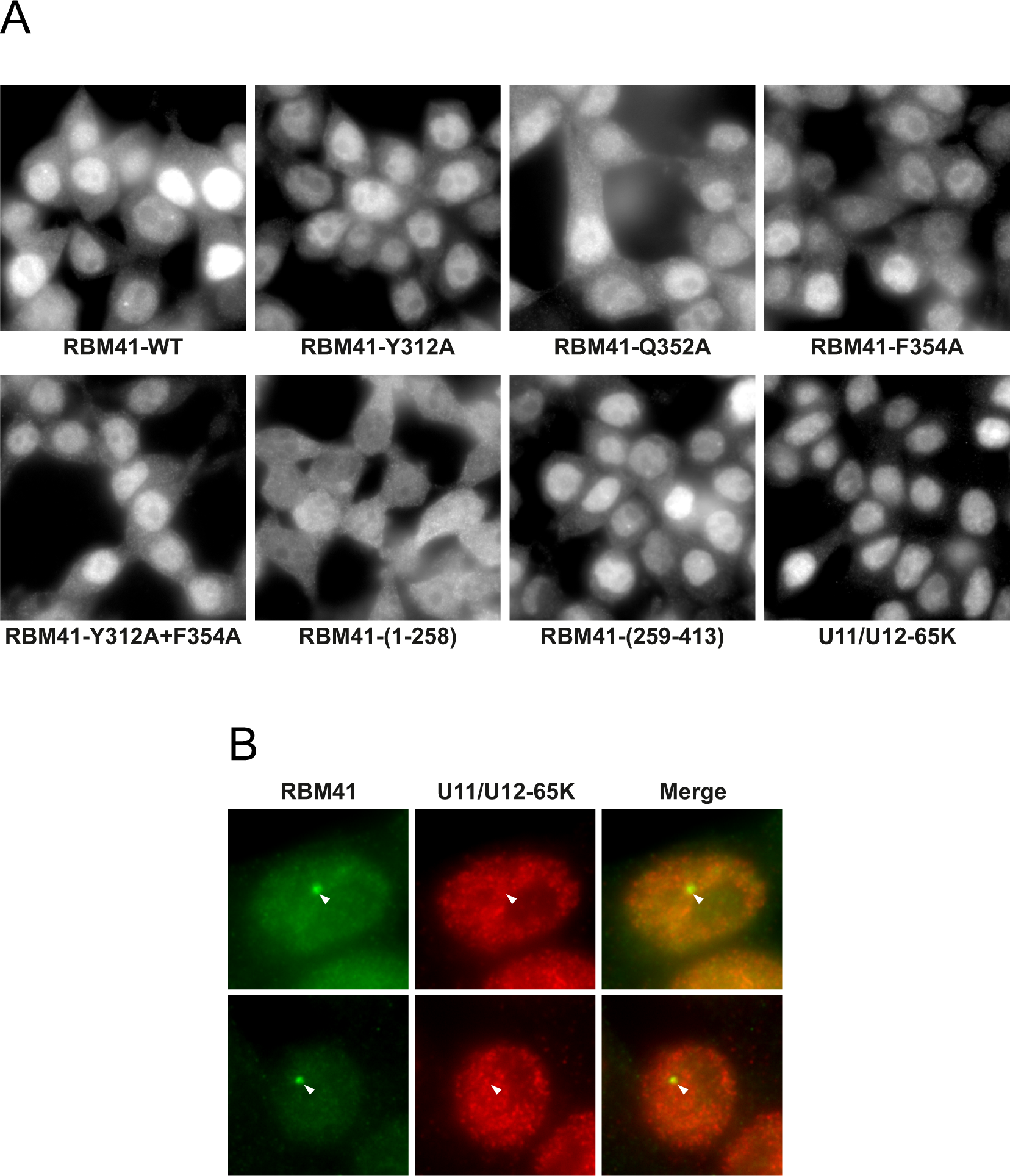
(A) Localization MAC-tagged constructs in Flp-in 293 cell lines. Expression was induced with tetracycline for 24 h and immunofluorescence carried out with anti-HA antibody. **(B)** Immunofluorescence staining of HEK293 cells for with anti-U11/U12-65K and anti-RBM41 antibodies.

**Supplementary Figure 5.**
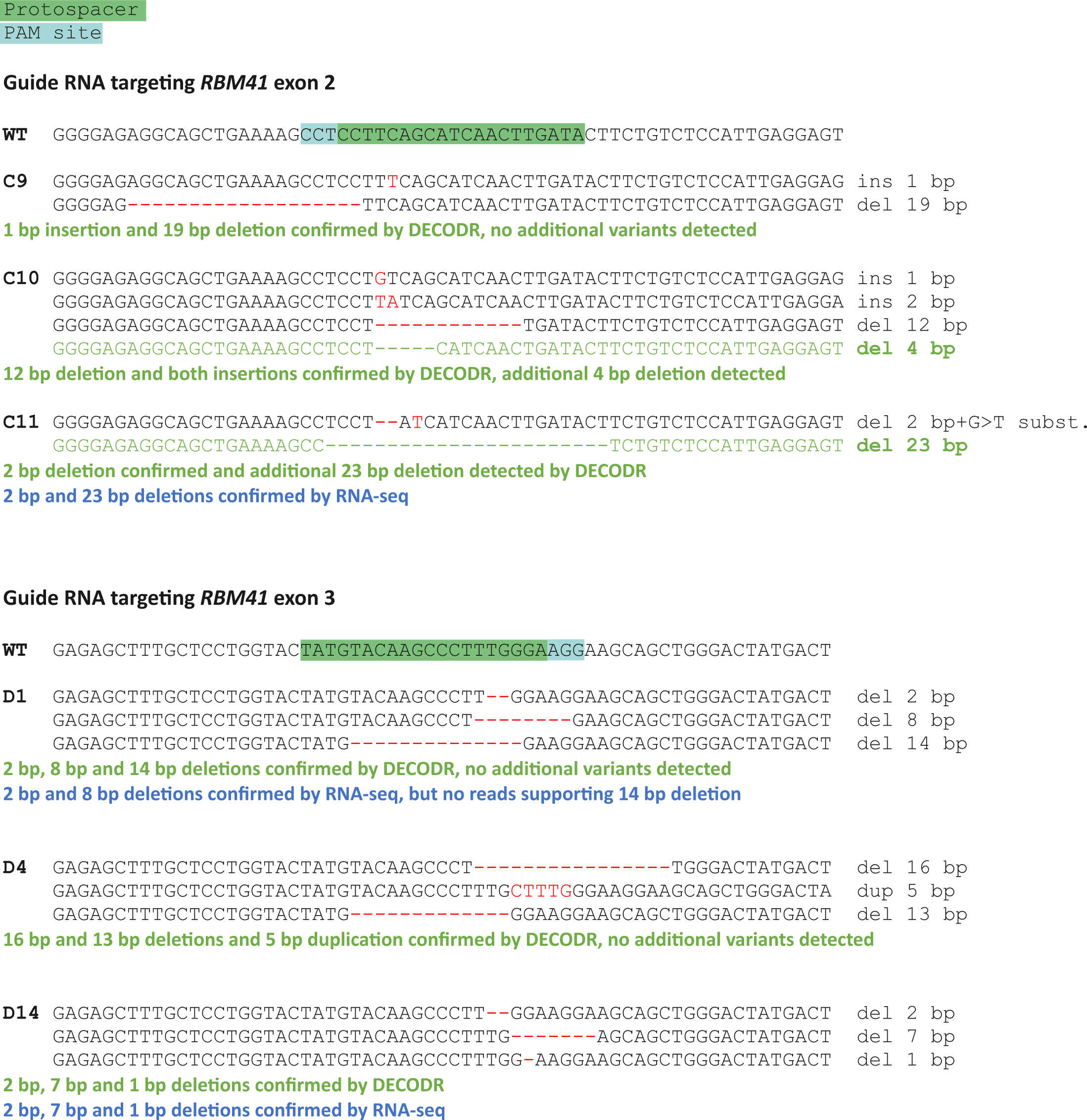
The sequeces of individual HEK293 the RBM41 Knock-Out clones. The targets of the Guide RNAs in RBM41 exons 2 and 3 are indicated in the WT sequece. Clones C11, D1, and D14 were used in the RNAseq analysis. C9, C10, and D4 were used in RT-PCR validation experiments.

